# ATR-I774Yfs*5 promotes genomic instability through micronuclei formation

**DOI:** 10.1101/2022.03.10.483800

**Authors:** Nathaniel Holcomb, Bithika Dhar, Hong Pu, Robert-Marlo Bautista, Anna Heink Overmann, Lauren Corum, Brent Shelton, John D’Orazio

## Abstract

Although mismatch repair (MMR) defects are associated with high risk of malignancy, the specific oncogenic drivers pertinent to MMR-affected cancers are poorly characterized. The heterozygous ATR-I774Yfs*5 mutation, the result of strand slippage in a poly-A tract of the Ataxia Telangiectasia and Rad3 related (ATR) gene, is overexpressed in MMR-defective malignancies including colorectal carcinoma (CRC) and is the most common ATR mutation in cancer. Here, we explore the contribution of ATR-I774Yfs*5 to genomic integrity. Using heterozygous ATR-I774Yfs*5 HCT-116 cells to mimic the native mutation, we found this mutation reduced ATR activity as measured by damage-induced Chk1 phosphorylation at S317 and ATR autophosphorylation ATR at T1989. ATR-I774Yfs*5 expression impaired genomic stability as visualized by the appearance of micronuclei in two stable expression models as well as in cell lines transfected with ATR-I774Yfs*5. Micronucleus development was dependent on replication and independent of ATR copy number. ATR-I774Yfs*5 expression did not alter cellular viability, cell cycle progression, or replicative rate, suggesting this mutation is well-tolerated despite its destabilizing effect on the genome. Taken together, these data suggest that the ATR-I774Yfs*5, whose development is favored in the context of MMR deficiency, may represent an important driver of a mutator phenotype by promoting genomic instability.

## 1 Introduction

Colorectal cancer (CRC) is one of the leading causes of cancer-related deaths in the US and worldwide, resulting in an estimated 53,200 deaths in the US in 2020 [1, 2]. A seminal report on microsatellite instability in CRCs separated cancers into two groups based on exonic mutational frequencies [3]. The higher mutational rate group, defined by a mutational frequency of higher than 12 per 10^6^, was described as “hypermutated” and had a very high incidence of microsatellite instability (MSI) that was mostly absent in the cancers with a lower mutational rate. MSI high cancers are typically characterized by deficiencies in the mismatch repair (MMR) pathway, a genome maintenance system charged with replicative fidelity by correcting mismatched bases or small insertions or deletions errantly inserted by replicative DNA polymerases [4]. MMR deficiencies can be inherited, such as those that occur in Lynch Syndrome, or they can be sporadic [5]. Without sufficient MMR, replicative fidelity is diminished and mutations accrue at an accelerated rate due to the inherent error rates of replicative polymerases [6, 7]. This “mutator phenotype” is thought to be an important contributor to carcinogenesis by accelerating genomic instability [8-11]. MMR is especially critical for repairing mispairs that occur during DNA replication of polynucleotide track repeats as replicative polymerases are prone to slipping at these nucleotide stretches [4]. Indeed, microsatellites are thought to occur predominantly in poly-adenosine (poly-A) tracts due to deletion of at least one adenine residue due to unrepaired strand slippage during DNA replication due to MMR deficiency [12].

Here we report how ATR-I774Yfs*5, a truncating mutation in the ATR gene overrepresented in MSI-high cancers, may represent an important oncogenic driver mutation by promoting genomic instability. ATR is a phosphatidylinositol 3-kinase related kinase that acts as an important DNA damage sensor and control molecule, particularly after cellular injury that results in tracts of single-stranded DNA [13]. It is also essential for replication fork stability as evidenced by the embryonic lethality of ATR-null mice as well as the double stranded DNA breaks, chromosome shattering and mitotic catastrophe that result from homozygous ATR loss [14, 15]. ATR-I774Yfs*5 is a frameshift mutation in a 10-A polyadenine repeat in exon 10 of ATR, and since polymerase strand slippage is normally recognized and repaired by MMR, the mutation occurs preferentially in MMR deficient cancers [16]. We found that stable expression of ATR-I774Yfs*5 impaired canonical ATR signaling, as measured by a reduction in p-Chk1 (S317) or p-ATR (T1989) after damage relative to WT/WT ATR expressing cells, suggesting that the mutation results in a dominant negative ATR physiology. Additionally, we observed that ATR-I774Yfs*5 induced DNA damage, as seen by the accumulation of micronuclei in the absence of exogenous damaging agents. Surprisingly, however, ATR-I774fs*5 did not impact cell viability, replicative rate or cell cycle progression, despite the well-characterized involvement of ATR in these processes. ATR-I774Yfs*5-mediated micronuclei induction was replication-dependent, confirming that the micronuclear phenotype relied on cell division. Together, these findings indicate that ATR-I774Yfs*5 expression may lead to genomic instability and a blunted DNA damage response, thereby contributing to oncogenesis.

## 2 Materials and Methods

### 2.1 *In silico* work

The *in-silico* data collected on ATR-I774Yfs*5 mutational frequency was obtained from publicly-available cBioPortal and COSMIC databases. We observed multiple names for the entry of this mutation, including I774Yfs*5, I774Nfs*3 (which represent a single A deletion or insertion in the poly-A tract, respectively) as well as mutation names such as “K771Yfs*5”, “I774Yfs*3”, “K773Nfs*3”, “P775Tfs*5”, and “I774Kfs*6”. We determined that these different mutational identifiers were all referring to a insertion or deletion in the 10-A stretch in ATR encompassing codons 771 to 774, and as such all encode a severely truncated ATR. While the insertion and deletion mutants generate slightly different protein products (the premature termination codon is 2 amino acids apart), they were analyzed together to create a profile of frequency and types of cancers harboring truncated ATR mutants created from the frameshift event at this polynucleotide region. The co-incidence of truncated ATR with MMR deficiencies was determined by probing the mutation profiles of all patients with truncated ATR for mutations in proteins contained in the MutSα and MutLα complexes.

### 2.2 Cells and cell culture

HCT-116 (ATCC CCL-247) and DLD-1 (ATCC CCL-221) human colorectal cancer cell lines, as well as NCM-356 (ATCC CVCL-D875) non-transformed human colonic mucosal cells were maintained in McCoy’s media (Thermo Scientific) supplemented with 10% FBS (Gemini Bio-Products). HIEC-6 (ATCC CRL-3266), a benign colonic epithelial cell line, was maintained in Opti-MEM (Thermo Scientific) supplemented with 20 mM HEPES, 10 mM Glutamine, 10 ng/mL EGF, and 4% FBS (all Thermo Scientific). All cells were maintained in a 37°C incubator with 5% CO_2_.

### 2.3 Stable expression of truncated ATR

HCT-116 cells were modified to express truncated ATR using two separate systems. First, a plasmid containing the truncated ATR cDNA (containing the first 779 codons only) and a Kanamycin resistance marker was obtained (System Biosciences, SBI, Palo Alto, CA). This plasmid was built into SBI’s FC500A-1 vector that contains attB sites which allow the target sequence to be stably integrated into mammalian genomes in low copy number when co-transfected with ϕC31 integrase) [17]. Following co-transfection, cells were grown in selective medium containing 400 μg/mL G418 to select for stable expression of the ATR plasmid. We also generated an HCT-116 GFP empty vector stable clone using the same process. Second, ATR heterozygous knock-in HCT-116 clones labeled as D15, M10, and M18 were generated (Synthego, Menlo Park, CA), with sequence validation of the mutation of one allele of ATR to I774Yfs*5 (See Supplemental Fig. 1 for confirmation of heterozygotic knockin), as well as a WT HCT-116 reference cell line.

### 2.4 Plasmids

ATR-WT-GFP is a modification of the PCDNA 3.1 ATR-WT plasmid (Addgene, Watertown, MA). A GFP tag was added by Applied Biological Materials (Richmond, BC). ATR-I774Yfs*5-GFP is a modification of the ATR-WT-GFP PCDNA 3.1 plasmid, with an additional modification of a deletion of 1 adenosine in the 10-A polynucleotide sequence in exon 10 (codons 771-774) of the ATR cDNA sequence (Applied Biological Materials).

### 2.5 Antibodies/Confocal Materials

The following antibodies were used in Western blotting and confocal microscopy experiments: Chk1, ATR, p-Chk1 (S-317), and Lamin A/C primary antibodies (Cell Signaling Technologies, Danvers, MA), and HRP-conjugated secondary antibodies; FITC conjugated Donkey anti Rabbit IgG and TRITC conjugated Donkey anti Mouse IgG (Jackson Immunoresearch, West Grove PA). Cell membrane boundaries were detected using Alexa Fluor 647 Phalloidin (Cell Signaling Technologies). Nuclear staining was detected using Prolong Diamond antifade DAPI mounting solution (Invitrogen, Waltham MA).

### 2.6 Transfections

Unless otherwise noted, transfections were performed 24 hours prior to phenotypic analysis or experimentation. HCT-116, DLD-1, and NCM-356 cells were seeded the day before transfection, and transfected using TurboFect (Fisher Scientific, Waltham, MA) according to manufacturer’s instructions.

### 2.7 Western Blot analysis

HCT-116 WT, clones D15, M10, and M18 cell lines were plated at 100,000 cells/well in 6-well plates, grown overnight and then treated ± 5 μM Doxorubicin for 24 hours (37°C, 5% CO_2_). After treatment, cells were collected in cold PBS by cell scraping, pelleted and washed, then lysed in 1xRIPA buffer (EMD Millipore Corporation, Burlington, MA) containing 1x Halt Protease and Phosphatase Inhibitor cocktail (Thermo Scientific) and 20 nM dithiothreitol (Thermo Scientific). Samples were sonicated using a Microson ultrasonic cell disruptor, protein concentrations were calculated using the Pierce TM BCA Protein Assay (Thermo Scientific). Samples were diluted in Nupage LDS 4x loading dye (Invitrogen), heated to 95°C (5 minutes), and run on precast mini-protean TGX gradient gels (Bio-Rad) according to standard procedures. After running, samples were transferred onto PVDF (Bio-Rad), blocked in TBS-T containing 5% milk powder, and incubated with the primary antibodies as indicated (1:1000 in TBS-T with 5% milk powder) overnight in 4°C. Following incubation, membranes were washed with TBS-T, incubated with the appropriate anti-mouse and anti-rabbit antibodies (1:10,000 in TBS-T with 5% milk powder) for 2 hours at RT. Following secondary antibody incubation, the membranes were washed in TBS-T and visualized using ECL+ (Thermo Scientific).

### 2.8 CFSE dye-dilution assay

HCT-116 WT, clones D15, M10, and M18, as well as HCT-116 K779* and HCT-116 GFP were plated at 100,000/well (early timepoints) and 25,000/well (later timepoints) to maintain cells in log growth phase throughout the experiment in 6-well plates and allowed to adhere overnight. The next day cells were fed 20 μM CFSE (Thermo Scientific), or mock treated for 10 minutes at 37°C, then medium was added to quench the feed. At 24, 48, 72, and 96 hours post-CFSE feed, one well of each condition was collected and fixed in 70% ethanol. After completion of the timecourse, samples were resuspended in 150 μL PBS for flow cytometric analysis of CFSE fluorescence using the Markey Cancer Center Flow Cytometry and Immune Monitoring Shared Resource Facility. A single value representing population doubling time was calculated using an exponential decay regression analysis of the CFSE signal from each of three independent experiments.

### 2.9 Annexin/PI viability assay

HCT-116 WT, clones D15, M10, and M18, as well as HCT-116 K779* and HCT-116 GFP were seeded in triplicate in 6-well plates at 100,000 cells/well and grown for 24 hours. Log-phase cells were trypsinized, pelleted, washed in PBS, pelleted again, then analyzed for viability using the FITC Annexin V Apoptosis Detection Kit (Thermo Scientific). Briefly, cells were resuspended in 1xAnnexin Binding buffer in PBS at 1×10^6^ cells/mL, 150 μL of that was transferred into a 5-mL polystyrene tube, and 5 μL each of FITC Annexin-V and propidium iodide (from the kit) were added to each sample. The samples were then analyzed by flow cytometry using the Markey Cancer Center’s Flow Cytometry and Immune Monitoring Shared Resource Facility. Values were reported as ± Annexin and ± PI relative to appropriate control samples, and data are presented as the average of Annexin/PI -/- cell populations from three independent experiments.

### 2.10 Confocal imaging

HCT-116, DLD-1, NCM-356, and HIEC-6 cells were seeded onto 8-well Lab-Tek glass chamber slides (Nunc, Rochester, NY) at 10,000 cells/well, grown to approximately 75% confluence, and transfected with ATR-I774 (wild type), ATR-I774Yfs*5, or mock transfected for 24 hours. Following transfection, cells were fixed with 4% paraformaldehyde, permeabilized with 0.5% Triton-X in PBS, blocked with 10% Donkey Serum, then incubated overnight in the desired primary antibodies (all at 1:100 in PBS). The slides were then washed in PBS and incubated with FITC conjugated Donkey-anti Rabbit IgG and TRITC conjugated Donkey-anti Mouse IgG fluorescently tagged secondary antibodies (1:100 in PBS) for two hours at RT. After incubation, slides were washed in PBS and incubated with Alexa Fluor 647 Phalloidin (1:50 in PBS) for 30 minutes. After a final wash in PBS, slides were mounted with a DAPI mounting solution, and visualized with a Nikon eclipse Ti2 confocal microscope with a Nikon LU-NV light source. HCT-116 K779* and HCT-116 CRISPR clones were seeded in the same manner and fixed after reaching the desired density, following the same steps as above.

### 2.11 Thymidine blockade

Cellular replication was inhibited by culturing cells in 2mM thymidine. After 24 hours, cells were fed 5 μM EdU, a thymidine analog, in the culture medium (± 2 mM Thymidine) for an additional 24h. EdU incorporation was quantified using an Alexa Fluor 594 Click-IT EdU detection kit (Thermo Scientific) and confocal microscopy from each of three independent experiments and presented as the presence or absence of EdU in cells and micronuclei in each of the four cell clones examined (HCT-116 WT, ATR mutant clones D15, M10, and M18).

### 2.12 Statistics

Statistical analysis was performed using GraphPad PRISM 5 software (GraphPad Software, San Diego, CA) as well as Sigma Plot 14 software (Systat Sotfware, San Jose, CA) and done in collaboration with Dr. Brent Shelton of the Markey Cancer Center’s Biostatistics and Bioinformatics Shared Resource Facility. Data are expressed as mean ± SD. For data with a mean and standard deviation, a 1-Way ANOVA (with appropriate post-tests) was used to determine the statistical significance of differences between groups; Fisher’s Exact test (SAS software, Cary, NC) was used to determine the statistical significance of differences when mean values were unobtainable (as in the case of MN incidence across samples with varying sample sizes per field). For all experiments, a p value ⩽ 0.05 was considered statistically significant.

## 3 Results

### 3.1 ATR-I774Yfs*5 is the most common ATR mutation across human malignancies

The ataxia telangiectasia and rad 3 related (ATR) gene encodes a phosphatidylinositol 3-kinase-related kinase (PIKK) that functions as a master regulator of DNA damage responses and that is essential for proper DNA replication by stabilizing replication forks (reviewed in [18]. We combed online databases for clinically relevant ATR mutations in cancer and found that of the over 1700 ATR mutations and 330 truncating mutations across human cancer types, ATR-I774Yfs*5 is the most common (Fig. 1A), being found in over a dozen different cancers and most commonly encountered in colorectal, endometrial, gastrointestinal, and pancreatic cancers (Fig. 1B). Over 32% of all the observed ATR-I774Yfs*5 mutations were found in colorectal cancers. The mutation occurs in the context of a run of 10 adenosines (Fig. 1C) that makes the region prone to polymerase strand slippage and subsequent frameshift mutations [12]. Depending on where the insertion or deletion event is assigned, some of the mutations are classified at position 771 (vs. 774), but any of the insertions/deletions in this poly-A run result in frameshifts that introduce a stop codon 3-5 amino acids downstream to yield a severely truncated ATR protein. Thus, we consider these mutations to be functionally equivalent. These mutations (I774Yfs*5, I774Nfs*3, I771Nfs*3, etc.) are especially common in the context of microsatellite instability [19, 20] presumably because DNA polymerase-mediated strand slippage events would be less likely to be repaired in a mismatch repair defective background [19, 20]. Indeed, there is a high co-incidence of the ATR-I774Yfs*5 mutation with mutations in the proteins comprising the MutSα and MutLα complexes of the MMR pathway, which are involved in recognizing and repairing mispairs and small insertion/deletion events [21],with co-incidence rates of 14-47% between ATR I774fs*5 and each protein in the complex (Fig. 1D). A truncated ATR protein resulting from expression of the ATR-I774Yfs*5 mutation would theoretically be able to bind to RPA and/or ATRIP and retain its nuclear localization sequence (NLS) as those domains are predicted to persist in the truncated protein. However, the mutant protein would be kinase-dead and lack several other important protein domains (Fig. 1E, F), raising the possibility that the truncated ATR protein product could function in a dominant-negative manner. Because of its frequency compared to other ATR mutations across cancer types, because it was found in colorectal carcinoma at a greater frequency than other cancers, and because of the dearth of information about this particular mutation, we investigated the impact of ATR-I774Yfs*5 on genomic stability in colorectal cell lines.

**Figure 1:**
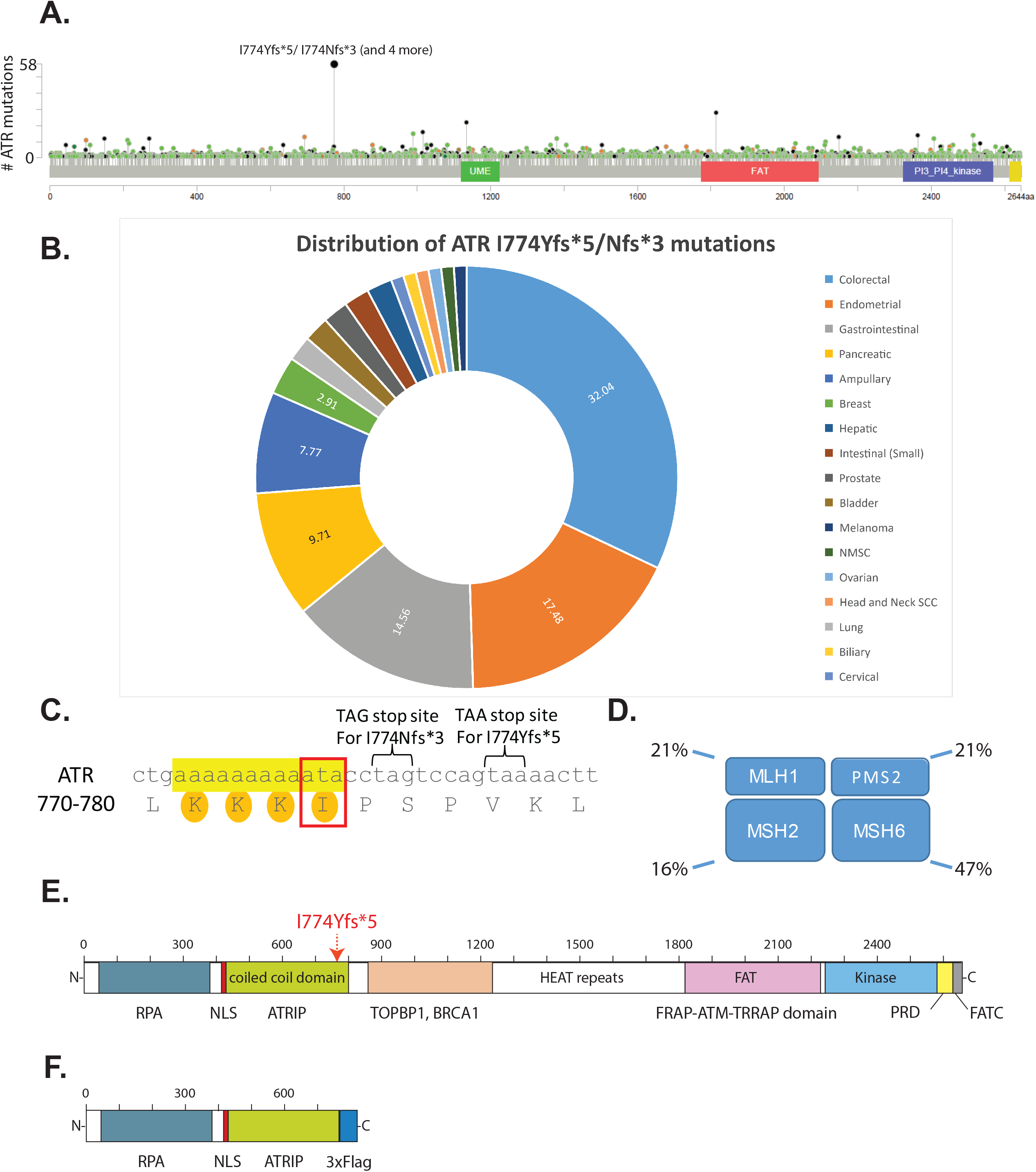
*In silico* analysis of ATR-I774Yfs*5/Nfs*3, **(A)** Lollipop plot of ATR mutations across all cancer types; there is a mutational hotspot at the codon 774 which occurs at a far greater frequency compared to any other ATR mutation. **(B)** Distribution pattern of all identified ATR I774Yfs*5/Nfs*3 mutations from publicly available databases; cancers typically associated with microsatellite instability (colorectal, endometrial, gastric, and pancreatic cancers) account for about two thirds of all identified ATR-I774Yfs*5/Nfs*3 mutations. **(C)** Nucleotide context of these truncation mutations show they are frameshifts that occurs as the result of a single adenosine insertion (I774Nfs*3) or deletion (I774Yfs*5) in a poly-adenosine region of ATR producing a truncation mutation at codon 776 (TAG stop codon) or codon 778 (TAA stop codon) for the insertion or deletion events, respectively. While the insertion or deletion of a single A could theoretically occur at any nucleotide along the poly-adenosine sequence, the resulting amino acid change only begins at codon 774 (boxed). **(D)** Co-incidence between all observed ATR-I774Yfs*5/I774Nfs*3 mutations and mutations in genes in the MutSα /MutLα complexes. Some ATR mutants were co-incident with multiple MutSα/MutLα complex genes, while some had no co-incidence. **(E)** A map of the functional domains of ATR, showing the location of the mutation of interest, ATR-I774Yfs*5 **(F)** ATR-I774Yfs*5/I774Nfs*3 mutations generate a putative truncated ATR product which lacks numerous functional domains including the C-terminal kinase domain but still retains the RPA and ATRIP interaction domains as well as the nuclear localization sequence (NLS).

### 3.2 Cells stably expressing truncated ATR are viable and do not have impacted replication or cell cycle progression

Cliby and colleagues reported that expression of a mutant ATR truncated at codon 767 inhibited ATR-dependent Chk1 phosphorylation in response to UV-mediated DNA damage in K562 human leukemia cells [19], suggesting dominant negative inhibition of native ATR by a truncated ATR protein fragment. We recognized that transient transfection of ATR-I774Yfs*5 does not replicate the biology of what must necessarily be heterozygous expression of this mutation, as both copies of WT-ATR are still present in the transfected cells and ectopic expression may lead to overexpression of the transgene. We therefore employed two systems for stable expression of truncated ATR into HCT-116 CRCs. First, we engineered a CRISPR-Cas9 mediated mutation of one allele of ATR to the ATR-I774Yfs*5 mutant in HCT-116 colorectal carcinoma cells to generate three unique ATR-heterozygous clones that mimic the native heterozygous mutation found in an MMR-defective background, identified as cell lines D15, M10, and M18. Next, to address the possibility that the resulting phenotype could be trivially explained by ATR haploinsufficiency, we generated a ϕC31 integrase mediated low-copy insertion of a truncated ATR cDNA into ATR-WT/WT HCT-116 cells. This ϕC31 integrase system approach yielded one HCT-116 cell line expressing ATR-K779* (a truncated cDNA insert to mimic the length of ATR-I774Yfs*5) as well as a GFP stable control line. Each of these stable truncated ATR-expressing cell lines were examined for viability, replicative proficiency, and cell cycle stability relative to their respective controls in the absence of exogenous injury. Using an Annexin/PI assay, we observed no significant difference in steady-state viability between ATR mutant cells and ATR WT cells in both systems (Fig. 2A). CRISPR ATR mutant cells had a viable population of between 88.2% and 90.9%, while WT ATR cells were 91.9% viable (*p* > 0.05 using a 1-way ANOVA). Viability for K779* and GFP Empty vector stable HCT-116 cells were 92.4% and 91.3%, respectively (*p* > 0.05 using a Student’s T-test).

**Figure 2:**
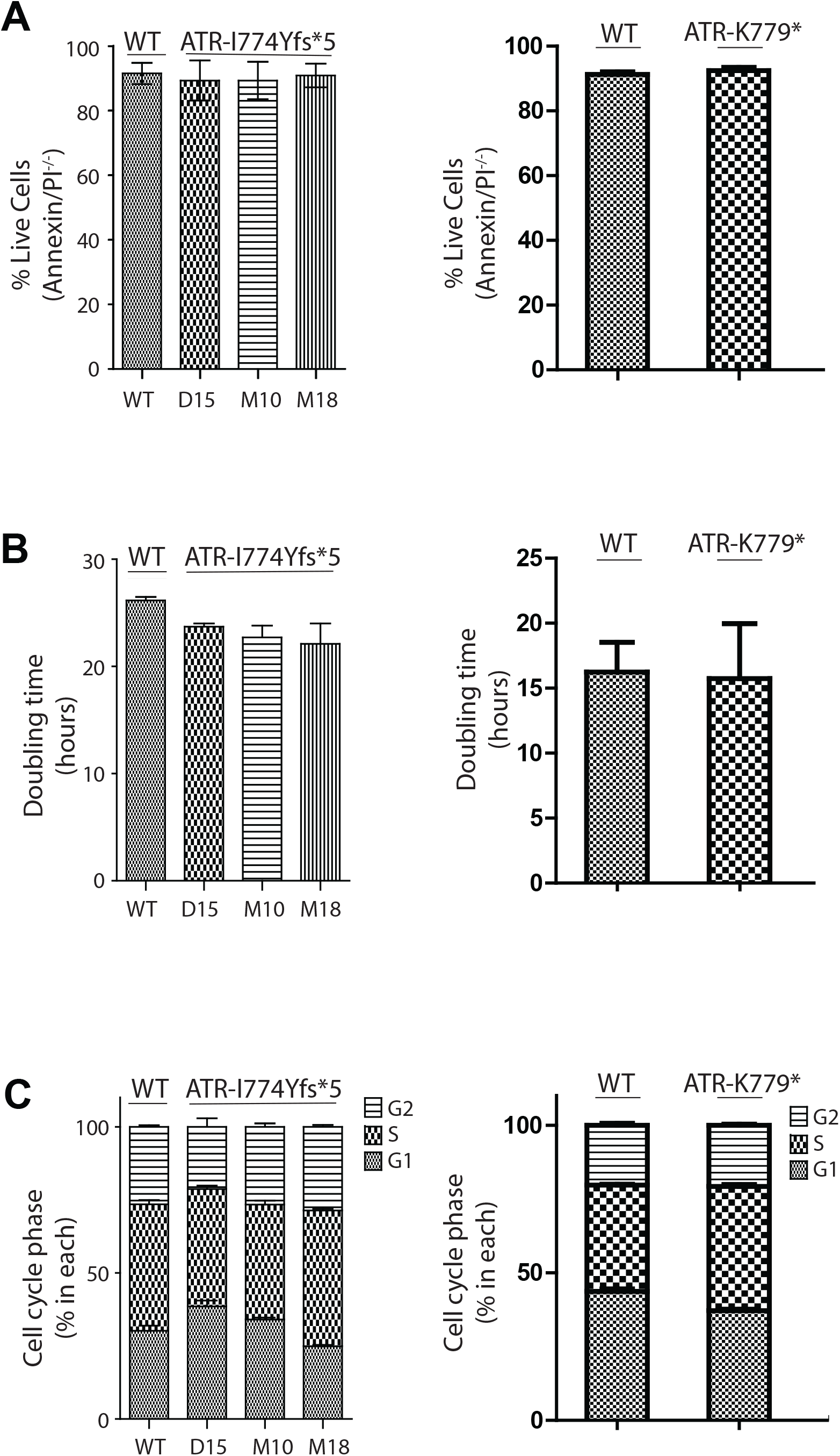
Introduction of the truncated ATR mutation does not cause apoptosis and does not impact cell doubling time or cell cycle. **(A)** Annexin-PI viability assays of CRISPR ATR mutants (left) and ϕC31 Integrase-mediated ATR mutants (right) show no significant reduction in viability relative to ATR-WT cells, as measured by flow cytometry (*p* > 0.05). **(B)** Cellular replicative rate, as measured by population doubling time using a CFSE dye-dilution assay, was unchanged in both CRISPR ATR mutants (left) and ϕC31 Integrase-mediated ATR mutants (right), relative to their ATR-WT counterparts (*p* > 0.05). **(C)** Steady-state cell cycle phase analysis was measured in the stable ATR mutants using PI and flow cytometry. In both the CRISPR ATR mutants (left) and ϕC31 Integrase-mediated ATR mutants (right), there was no significant change in percentage of cells in any phase of the cycle relative to ATR-WT counterparts (*p* >0.05). Data for **A-C** are from three independent experiments and are presented as mean **±** standard deviation of those experiments.

We also evaluated the impact of truncated ATR on cellular replicative rate using a CFSE dye dilution assay. Although we anticipated that incorporation of a mutant truncated ATR with potential dominant negative physiology might impair replication and therefore prolong cell division, in fact we observed a trend in the opposite direction (Fig. 2B). Thus, for the CRISPR-derived lines, the WT control HCT116 cells (with two wild type copies of ATR) exhibited a mean doubling time of 26.2 ± 0.52 hours while D15, M10 and M18 CRISPR ATR mutant lines had faster average doubling times of 23.71± 0.53, 22.71 ± 1.92, and 22.12 ± 3.30 respectively. Through three experimental replicates, however, the differences did not reach statistical significance (*p*=0.13) as measured by one-way ANOVA analysis. Similarly, for the ϕC31 integrase system, ATR-WT control HCT-116 cells had an average doubling time of 16.0 ± 4.2 hours, and their ATR-K779* integrated passage-matched counterparts exhibited a mean doubling time of 15.3 ± 2.8 hours (*p* > 0.05). Thus, we concluded that stable expression of the truncating ATR mutations tested did not affect cell doubling time.

Lastly, we explored whether the truncating ATR mutation might impact cell cycle. When comparing D15, M10, and M18 HCT-116 CRISPR ATR mutant lines to WT control HCT-116 cells, there were no statistically significant differences in percentages of cells in G_1_ (*p*=0.19), S (*p*=0.17), or G_2_ (*p*=.021) phases as measured by one-way ANOVA. (Fig. 2C, left). Similarly, the K779*-expressing cells showed no statistically significant difference in percentage of cells in G_1_ (*p*=0.90), S (*p*=0.95), or G_2_ (*p*=0.73) phases relative to their control cells, as measured by a Student’s t-test (Fig. 2C, right). Based on these findings, we concluded that stable expression of either heterozygous ATR-I774Yfs*5 or of low copy number K779* did not have a reproducible impact on cell cycle. As the CRISPR-generated cells are true heterozygotes and therefore more closely mimic malignancies naturally expressing the mutation, they were used for the majority of subsequent analyses.

### 3.3 Truncated ATR suppresses endogenous ATR activity in colorectal cells

In order to determine the functional impact of ATR-I774Yfs*5 in ATR-mediated damage signaling, we examined canonical ATR signaling in response to doxorubicin-induced DNA damage in the presence of heterozygous ATR-I774Yfs*5 vs homozygous ATR-WT. We exposed the ATR heterozygote CRISPR lines D15, M10, and M18, along with ATR-WT cells to 5 μM doxorubicin (or vehicle control) for 24 hours and examined multiple DNA damage responsive events, including Chk1 phosphorylation at Ser317, an ATR-dependent response to doxorubicin treatment [22] by Western blotting. As predicted based on heterozygous expression of a loss-of-function ATR-I774Yfs*5 mutant, ATR expression was reduced in the undamaged CRISPR mutant cell lines compared to ATR-WT cells, and the difference was significant in clones M10 and M18 (P < 0.05) (Fig 3A, B). After damage, ATR expression was similar between the ATR-WT and CRISPR mutant cell lines (p>0.05). We observed no differences between basal expression of total Chk1 in ATR-mutant cells compared to ATR-WT cells, and each of the cell lines exhibited a damage dependent decline in total Chk1 levels irrespective of ATR status [23] (Fig. 3A, C). Whereas ATR-WT cells showed a robust damage-induced increase in p-Chk1 (S317) in response to doxorubicin exposure (Fig. 3A, D), each of the ATR mutant clones demonstrated a blunted induction of p-Chk1 (S317) of only 37-43% as compared to WT (*p*<0.05). We also looked at p-ATR (T1989), a DNA-damage driven autophosphorylation event [24]. While WT-ATR expressing cells showed an increase in p-ATR (T1989) after doxorubicin exposure, this induction was substantially ablated in the mutant cells (Fig 3A, E) (*p* <0.05), suggesting another element of ATR signaling is negatively impacted in the ATR heterozygotes. From these data we conclude that the presence of the ATR-I774Yfs*5 mutation expressed in a heterozygous manner has no impact on total Chk1 levels but attenuates damage-induced phosphorylation of Chk1 on Ser317 and autophosphorylation of WT-ATR at T1989. Moreover, heterozygous ATR-I774Yfs*5 expression has mixed effects on total ATR expression in our cell lines.

**Figure 3:**
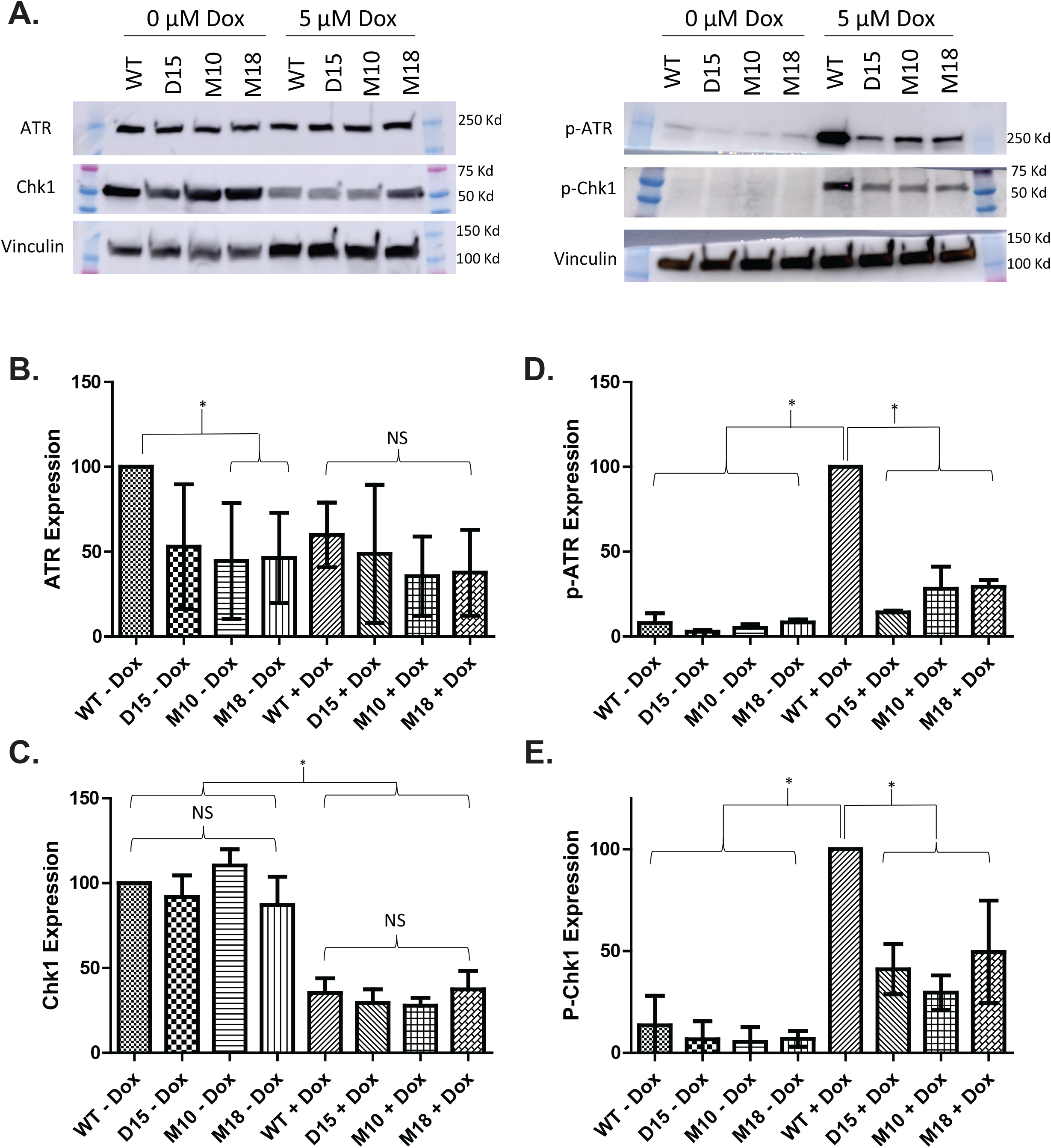
ATR-I774Yfs*5 exerts a dominant negative effect on ATR signaling after DNA damage. **(A)** Log-phase ATR heterozygotic mutant HCT-116 cell clones D15, M10, and M18 along with HCT-116 WT-ATR were treated with or without 5 μM doxorubicin for 24 hours. After exposure, cells were harvested and analyzed by Western blotting. Expression levels of ATR, Chk-1, pChk-1 (S317), and p-ATR (T1989) were measured using vinculin as a loading control. **(B, C, D, E)** Densitometry analysis across a minimum of 3 independent experimental immunoblot replicates (mean ± standard deviation) of total ATR expression (**B**), Chk-1 expression (**C**), p-ATR (T1989) expression (**D**) or p-Chk1 (S317) (**E**). For each graph, * denotes *p* <0.05 and NS indicates no statistical differences between groups compared using a 1-way ANOVA.

We also examined the impact of truncated ATR on p-Chk1 (S317) expression following doxorubicin exposure using confocal microscopy. In agreement with the immunoblot results, we observed that p-Chk1 (S317) expression was significantly induced in ATR-WT cells after doxorubicin exposure (Supplemental Fig. 2A, column 1). However, in the ATR-mutant expressing CRISPR cell lines this induction was attenuated (Supplemental Fig. 2A columns 2-4). Whereas doxorubicin induced a 256% increase in nuclear p-Chk1 (S317) in ATR-WT cells relative to undamaged controls (*p*< 0.05) each of the 3 CRISPR heterozygous ATR mutant cell lines exhibited attenuated p-Chk1 (S317) induction (D15; 71.0%, M10; 63.6%, M18; 29.4%; Fig 3E, F). While p-Chk1 (S317) induction after doxorubicin exposure was significant for each clone (*p* < 0.05), it was also significantly less than the induction seen in ATR-WT cells (*p* <0.05). Additionally, there was also no significant difference in basal p-chk1 expression levels in undamaged cells between ATR-WT and all three of the mutant cell lines (Supplemental Fig. 2B). Together, these data suggest that HCT-116 cells expressing ATR-I774Yfs*5 heterozygously exhibit an attenuated ATR-mediated DNA damage response.

Additionally, we examined the ability of ATR-I774Yfs*5 to impair ATR signaling in a homozygous ATR-WT background through transient transfection. We transfected full-length ATR-I774 (Wild-type, WT) and ATR-I774Yfs*5 plasmid constructs into HCT116 cells (24 hours) and subsequently exposed them to 10 μM doxorubicin (or vehicle control) for 1 hour (Supplemental Fig 2C). Doxorubicin exposure significantly increased p-Chk1 (S317) in both ATR-WT and ATR-I774Yfs*5 transfected cells (*p* < 0.05 for both comparisons), but the level of p-chk1 induction was attenuated by roughly 75% in ATR-I774Yfs*5 transfected cells relative to ATR-WT cells (Supplemental Fig. 2C). While ATR-WT cells increased p-Chk1 (S317) expression after doxorubicin treatment by 90%, the induction level in the ATR-I774Yfs*5 transfected cells was only 23% (Supplemental Fig 2D). Based on these findings, we concluded that ectopic transfection of ATR-I774Yfs*5 could phenocopy the negative regulation of ATR-mediated, DNA damage driven p-chk1 (S317) induction.

### 3.4 Expression of ATR-I774Yfs*5 is associated with the development of abundant micronuclei (MN) in CRCs

When examining confocal images of the ATR-mutant cell lines, we observed an increased incidence of micronuclei – defined as DAPI-positive signal outside the nuclear envelope but contained within the plasma membrane - in either the CRISPR-generated ATR-I774Yfs*5 heterozygous cells (Fig 4A) or the ϕC31-generated ATR-I779*-expressing (Fig 4B) HCT-116 cells. In both model systems, ATR mutant cell lines showed increased levels of micronuclei compared to their passage-matched ATR-WT control counterparts. The number of micronuclei in each expression system was significantly greater in the ATR mutant expressing cells than their WT counterparts (*p*<0.05 for HCT-116 GFP vs HCT-116 K779* and *p*<0.05 each for ATR WT vs D15, M10, and M18). Micronuclei are conventionally thought to be the product of clastogenic injury to DNA that results in strand beaks that could arise from aberrant replication, endogenous or exogenous DNA damage, or genomic instability (reviewed in [25]), thus we were interested in understanding this potential marker of genomic damage further.

**Figure 4:**
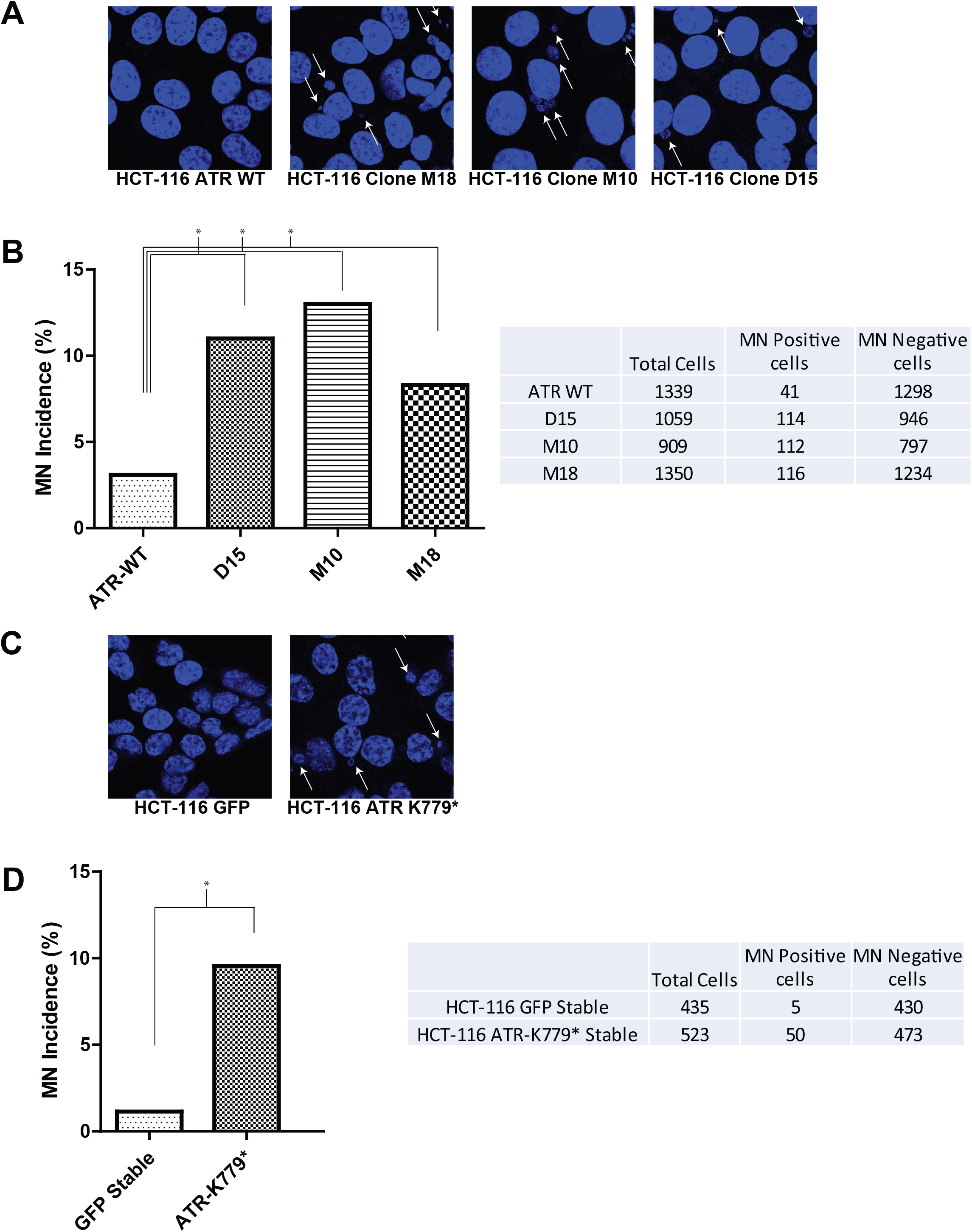
Stable introduction of ATR-I774Yfs*5 generates a micronuclear phenotype. **(A)** Representative images of log-growth phase HCT-116 cells with either WT ATR or CRISPR-engineered heterozygous ATR-I77fYfs*5 (lines M18, M10 and D15) DAPI stained to detect DNA; white arrows indicate micronuclei. (B) Quantification of micronuclei levels in HCT-116 cells with either WT ATR or CRISPR-engineered heterozygous ATR-I77fYfs*5 (lines M18, M10 and D15). DAPI staining outside the nucleus was used to define a micronucleus. D15, M10, and M18 all had significantly higher rates of MN formation compared to ATR-WT cells; * denotes statistical significance (*p* <0.05) for the indicated pairings using Fisher’s Exact test) **(C)**. Representative images of log-growth phase HCT-116 cells with either ATR-WT or ATR-K779* generated using the ϕC31 Integrase system. DAPI stained to detect DNA; white arrows indicate micronuclei. (D) Quantification of micronuclei levels in HCT-116 cells with either ATR-WT or ATR K779*. For each graph, * denotes statistical significance (*p* <0.05) using Fisher’s Exact test.

In an effort to determine whether the micronuclei phenotype was a consequence of the mutant ATR protein and not a byproduct of generating ATR-hypomorphic stable mutant cell lines (e.g. in the heterozygous CRISPR HCT116 system), we ectopically expressed ATR-I774Yfs*5 as well as ATR-WT across a panel of benign (HIEC-6, NCM356) and transformed (HCT116, DLD-1) colorectal cell lines, each harboring two wild type ATR alleles. In each cell line examined, transient transfection with ATR-I774Yfs*5 induced the development of micronuclei relative to ATR-WT transfected or mock-transfected cells (Supplemental Fig. 3A-B), suggesting that the phenotype is not simply due to reduced wild type gene dosage and that the phenotype is independent of MMR status. Notably, we observed marked differences in the morphology of micronuclei between cells harboring heterozygous (CRISPR) or low-copy number expression of truncated ATR vs. those transiently transfected with ATR-I774Yfs*5 plasmid. While the majority of micronuclei in CRISPR ATR mutant cells were surrounded by a laminin-containing envelope (Supplemental Fig. 4A, left) as is the case for conventional micronuclei [26], transiently transfected cells produce large quantities of extranuclear DNA that were not enclosed in lamin envelopes. We estimate that only approximately 15% of micronuclei had lamin containing envelopes in transiently ATR-I774Yfs*5-expressing cells (Supplemental Fig. 4A, right). In contrast, the majority (90-96%) of micronuclei in CRISPR ATR stable mutant cells were surrounded by lamin-containing envelopes (Supplemental Fig 4B). Additionally, transient transfection resulted in a significant number of extranuclear DNA fragments per cell, while stable expression of ATR-I774Yfs*5 generated MN at an incidence of typically 1-2 per cell (Compare Fig. 4 with Supplemental Fig. 3). We conclude that the introduction of truncated ATR either by transient transfection or by more physiologic low-copy number expression systems leads to micronuclei development and hypothesize that this phenotype is due to a dominant negative inhibition of WT ATR function.

### 3.5 ATR-I774Yfs*5-induced micronuclei are replication-dependent

We next explored the impact of replication arrest on the MN phenotype produced by ATR-I774Yfs*5. HCT-116 cells expressing ATR-WT or heterozygous ATR I774Yfs*5 via CRISPR knockin were fed 2 uM thymidine to arrest replication [27, 28]. After 24 hours in 2 mM thymidine (or vehicle control), cells were fed 5μM EdU for 24 hours in the presence of 2mM thymidine (or vehicle control). Following this, we examined EdU incorporation and micronuclei formation in the cells by confocal microscopy. Nuclear EdU incorporation was between 95 and 98% for ATR-WT and ATR mutant cells in the absence of thymidine (Fig 5A, C), indicating robust cell division in replication-proficient cells. In contrast, EdU incorporation was fully inhibited in the presence of the thymidine block in either ATR WT or ATR mutant cells. (Fig 5B, C), suggesting that thymidine incubation effectively halted cell division (*p*<0.05 comparing cellular EdU incorporation for all cell lines ± thymidine treatment using Fisher’s Exact test). We also observed that micronuclei in ATR-WT as well as ATR mutant cells also incorporated EdU at a high rate that was completely eliminated by thymidine treatment (Fig 5D), suggesting that MN development is replication dependent in both ATR-WT and ATR mutant cells (*p*<0.05 comparing EdU incorporation in MN for all cell lines ± thymidine treatment using Fisher’s Exact test). Nonetheless, some micronuclei were observed in replication-arrested lines, albeit at reduced levels in either ATR-WT or ATR mutant cells in the presence of thymidine-induced replication arrest compared to replication proficient cells (Fig. 5E). Specifically, WT-ATR MN incidence decreased from 4.4% to 1.8% after thymidine treatment, D15 MN incidence decreased from 10.8% to 4.1%, M10 MN incidence decreased from 14.1% to 8.5%, and M18 MN incidence decreased from 9.2% to 5.7% (*p*<0.05 for all comparisons using Fisher’s Exact test). We interpret these findings to suggest that halting replication prevented the formation of new MN, but did not eliminate the existing MN in these cells. Since the remaining MN were all EdU negative, we posit that they were formed prior to the introduction of the thymidine treatment and remained intact after replication was arrested (data summarized in Figure 5F). Together these data suggest that truncated ATR-dependent micronuclei formation is replication dependent and that micronuclei from prior divisions can be retained in cells.

**Figure 5:**
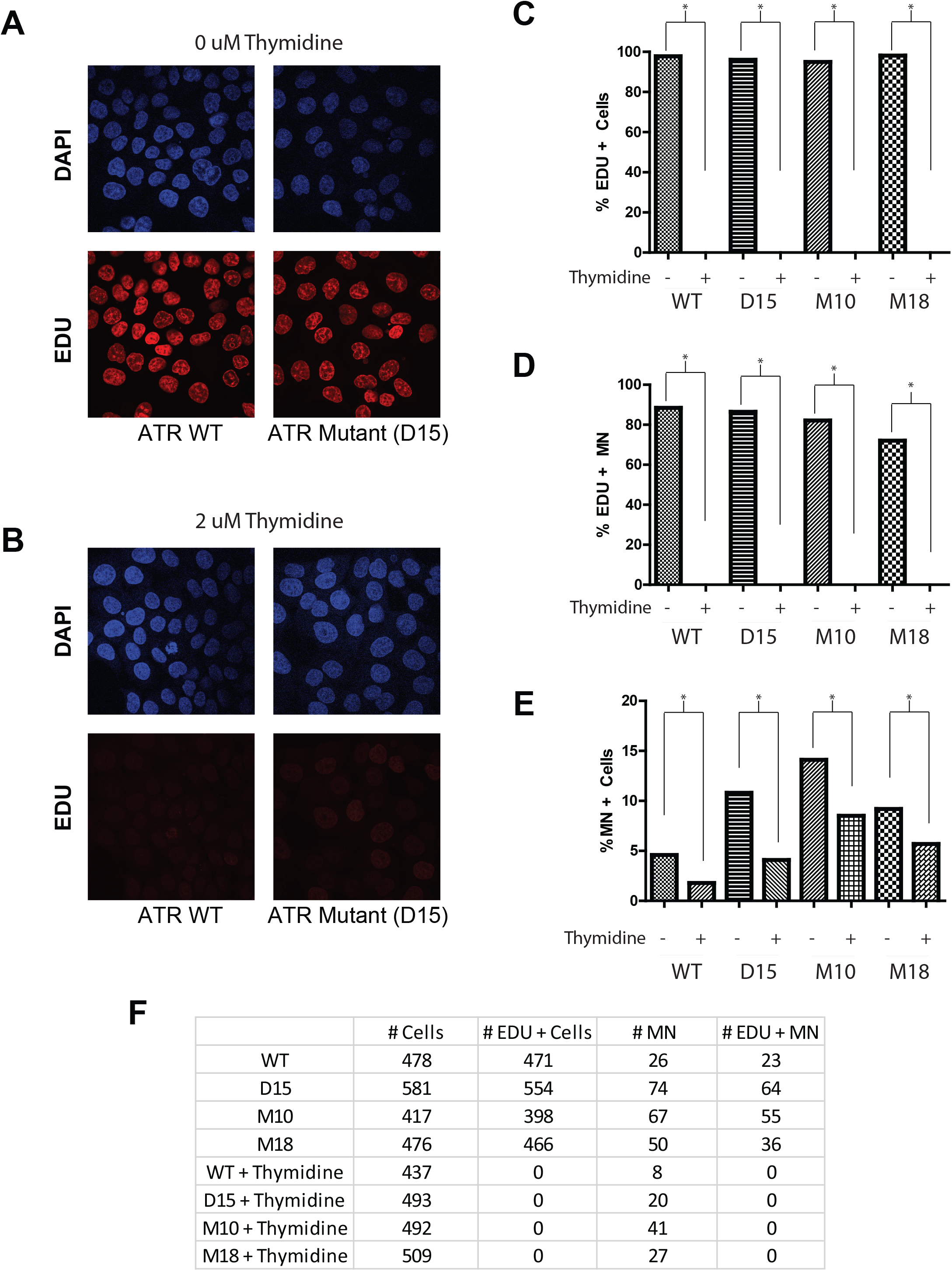
Micronuclei produced by heterozygous integration of ATR-I774Yfs*5 are replication-dependent. **(A)** representative images of log growth phase HCT-116 ATR-WT and D15 (heterozygous ATR-I774Yfs*5) labeled with EdU to show proliferation in the absence of thymidine treatment as indicated by red staining (DAPI staining to indicate DNA). **(B)** EdU incorporation into HCT-116 ATR-WT and a representative ATR heterozygotic mutant HCT-116 cell clone (D15) are shown in cells treated with 2mM thymidine for 24 hours prior to EdU treatment. **(C)** Quantification of EdU incorporation into nuclei of HCT-116 ATR-WT and all three ATR heterozygotic mutant HCT-116 cell clones ± 2 μM thymidine. **(D)** Quantification of EdU incorporation into observed micronuclei in HCT-116 ATR-WT and all three ATR heterozygotic mutant HCT-116 cell clones ± 2 μM thymidine. **(E)** Quantification of micronuclear incidence in HCT-116 ATR-WT and all three ATR heterozygotic mutant HCT-116 cell clones ± 2 μM thymidine. For **(C), (D)**, and **(E)**, * denotes *p*<0.05 using Fisher’s Exact test). **(F)** A chart depicting the cell counts for each clone in the two treatment conditions, along with EdU positive cell counts, total number of observed micronuclei, and the number of EdU positive micronuclei. These values were used to determine statistical significances as shown in (C), (D), and (E).

## 4 Discussion

In this study, we explored the impact of expression of the ATR-I774Yfs*5 mutation on colorectal cell physiology. We found that the ATR-I774Yfs*5 mutation destabilizes genomic integrity as evidenced by the generation of micronuclei. Importantly, the micronuclei generated by truncated ATR occurred independently of exogenous damage. We observed that ATR-I774Yfs*5 did not impact cell viability or replicative rate, but that replicative proficiency was required for micronuclear development. We envision that heterozygous presence of the ATR-I774Yfs*5 mutation impacts cell physiology by diminishing native ATR function enough to destabilize genomic integrity without leading to mitotic catastrophe which is typical when ATR dosage is too low, as is the case in a homozygous ATR-null state [29]. Since introduction of ATR-I774Yfs*5 did not impact cell viability or replicative rate, we conclude that the genomic instability caused by the mutation is well-tolerated by the cell. Together, these findings provide strong evidence to support ATR-I774Yfs*5 as a pro-mutagenic, DNA damage-inducing mutation that may have clinical significance in the genesis of several types of cancers, particularly MMR deficient, MSI high cancers where its incidence is favored.

A critical component in carcinogenesis is the accumulation of mutations which often inactivate tumor suppressors or activate oncogenes that fuel malignant degeneration. Indeed, while many mutations in key DNA damage/response genes may not be tolerated and lead to premature cell death, clonal evolution to a cellular background which promotes mutagenesis without enhancing apoptosis can ultimately lead to carcinogenesis because mutation-accumulating cells persist. The “mutator phenotype” found in MSI colorectal cancers is one clear example, and it is largely due to the fact that MMR is defective [9]. Miquel and colleagues reported that MMR-defective replicative infidelity leads to accelerated mutagenesis, especially at regions of polynucleotide repeats [16]. Specifically, they found that the 10-A polynucleotide tract in ATR (encompassing codons 771-774) was mutated (via a frameshift of one nucleotide) in 44% of patients examined in their cohort, a considerably higher incidence than the observed mutational frequency of ATR-I774Yfs*5 across cancers seen in the TCGA database. An additional study of this same ATR mutation found no mutations (0 out of 21) in MSS cancers but 33% of MSI CRCs (9 out of 27) contained the ATR mutation [30]. These studies suggest that while ATR mutations in general may not be very prevalent in cancers (about 4% according to the TCGA database), the ATR-I774Yfs*5 mutation appears to be overexpressed in certain populations, such as those individuals with impaired MMR and with resultant MSI tumors.

Importantly, our findings may not be limited to colorectal cancers; this same mutation was found to be overexpressed in MSI-high gastric tumors [31] as well as MSI-high endometrial cancers [20]. What remained unclear from prior studies, however, is whether this mutation drives carcinogenesis or is instead a passenger mutation. Our findings suggest that truncating ATR mutations may be active drivers of oncogenesis; however, further studies on oncogenic potential of ATR-I774Yfs*5 in non-transformed cells or in cells derived from other tissues will be required to truly confirm this mutation as an oncogenic driver. Critically, the mutation generates damage without impacting viability or replicative rate, perhaps due to the dominant negative impact ATR-I774Yfs*5 appears to have on wild-type ATR function. Thus, we posit that cells harboring the ATR-I774Yfs*5 mutation tolerate the levels of DNA damage caused by ATR-I744Yfs*5, resulting in the observed increase in micronuclei.

Previous reports suggest that partial loss of ATR in MMR-deficient, but not MMR-proficient, colorectal cancer cells generates DNA damage and micronuclei [32] as well as chromosomal instability [33]. Thus, ATR haploinsufficiency promotes genomic damage in certain conditions. At first glance, this could explain our observations, as the CRISPR-mediated ATR mutant cells were produced in HCT-116, an MMR-deficient cell line. However, both the ϕC31 Integrase-mediated ATR K779* cell line and all experiments involving ectopic expression of ATR I774Yfs*5 via transient transfection approaches generated MN on an ATR WT/WT background, suggesting that depletion of WT-ATR is not likely the sole explanation for the development of the observed MN phenotype. Indeed, our own data suggest that heterozygous ATR-I774Yfs*5 expression variably reduces total ATR levels (Fig. 3D). Given this, we infer that the observed micronuclear phenotype in cells expressing ATR-I774Yfs*5 is likely the result of the truncated ATR protein operating in a dominant negative manner to reduce cellular ATR activity to a level that interferes with its normal functions, resulting in impaired ATR signaling and ultimately micronuclei. Further studies of ATR-I774yfs*5 function in MMR proficient cell lines would help delineate the observed MN phenotype from that seen in ATR-depleted, MMR deficient cells.

Transient transfection with ATR-I774Yfs*5 into the panel of CRCs led to a profound micronuclear phenotype, but the morphology of these micronuclei did not match traditional micronuclei, which are typically few in number (per cell) and bound by an envelope containing lamin-A levels comparable to the parental nucleus [26]. The majority of extranuclear DNA observed in transiently transfected cells were not contained within laminar envelopes (Supplemental Fig. 4). In contrast, the great majority of extranuclear DNA observed in cells that stably expressed ATR-I774Yfs*5 were contained within laminar envelopes (Supplemental Fig. 4), identifying them as traditional micronuclei. The distinction is important as the CRISPR derived stable cells more appropriately mimic an endogenous heterozygotic ATR mutation event, and knowing that the consequence of this event is a traditional micronuclear phenotype allows us to examine the long-term consequences of these micronuclei. It is entirely likely that some of the micronuclei, presuming they are not toxic to the cell, will eventually be ejected into the extracellular space, as this is one of the fates of traditional micronuclei [34]. If this were to occur, this could possibly contribute to tumor progression through effects on cellular microenvironment such as initiation of extracellular DNA trap formation to promote inflammation [35]. Extracellular DNA has also been reported to promote tumor survival after chemotherapy [36] and even transformation of non-tumor cells [37]. Thus, cancers bearing an extra-cellular DNA shedding phenotype may be more aggressive than others, suggesting that ATR-I774Yfs*5 may be a biomarker of disease progression. Indeed, a previous work indicated that ATR-I774Yfs*5 was associated with reduced disease-free and overall survival in a small study of endometrial cancers [20]. We have not discovered the fate of micronuclei induced by ATR-I774Yfs*5, therefore more work needs to be done to understand the implications of this mutation on tumor progression.

It is important to note that despite our reproducible findings that micronuclei and reduced ATR function occur after stable introduction of the ATR-I774Yfs*5 mutation, we have not presented data actually documenting a truncated ATR protein in this setting. In fact, it is possible that NMD, a protective mechanism to protect the cell against damaging protein products arising from frameshift mutations introducing premature stop codons (reviewed in [38]), may process and degrade truncated ATR message [38]. It is possible that the CRISPR-generated cells express truncated ATR message (and protein) at a much lower frequencies than the ATR-WT message, which could explain why not all cells harboring ATR-I774Yfs*5 stably express MN, and perhaps why the mutation does not lead to the death of cells (by more complete ATR loss). While we found that full-length ATR expression is reduced in ATR-I774Yfs*5-containing cells (Fig. 3D), we were unable to detect truncated ATR protein by immunoblotting, perhaps due in part to the dearth of commercially-available ATR antibodies targeting the N-terminus of the protein, and our reliance on C-terminal-tagged stable ATR mutants which may lose their tag if the truncated protein is inherently unstable. We posit that if NMD is targeting the mutant mRNA, there may be a very low level of truncated ATR expressed– a level sufficient to induce the micronuclear phenotype but below detection limits of conventional immunoblotting.

Our observations support the concept that ATR-I774Yfs*5 is a genome-destabilizing mutation through dominant-negative ATR physiology, raising possibilities of novel therapeutic opportunities in ATR-I774Yfs*5-expressing cancers. Since loss of ATR below a certain threshold is not tolerated because of replicative catastrophe, it is possible that such tumors would be selectively sensitive to pharmacologic ATR inhibition or to inhibition of other DNA damage/response pathways (e.g. ATM), or indeed inhibitors of NMD (to increase truncated ATR levels to further inhibit ATR in cells). Future studies are planned to explore synthetic lethality possibilities.

## Abbreviations

MMR: (mismatch repair)
ATR: (Ataxia Telangiectasia and Rad3 related)
CRC: (colorectal carcinoma)
CFSE: (carboxyfluorescein succinimidyl ester)
EdU: (5-ethynyl-2’-deoxyuridine)
NMD: (nonsense-mediated decay)

## Acknowledgements

This work was supported by the following funding mechanisms: The National Institutes of Health [R01 CA131075], T32 CA165990, and the University of Kentucky [P30 CA177558]. This research was supported by the Flow Cytometry and Immune Monitoring, the Biostatistics and Bioinformatics and the Cancer Research Informatics Shared Resource(s) of the University of Kentucky Markey Cancer Center [P30CA177558]. We thank the Imaging Core facility of the COBRE [P20 GM121327] at the University of Kentucky for providing service with microscopes and imaging systems. Lastly, we acknowledge critical suggestions and input from Drs. David Orren, Katerina Zaytseva, Eva Goellner, Chi Wang, Daheng He, Jakub Famulski, Tianyan Gao and Christine Brainson.

