## Supplemental Fig 1-4 for "ATR-I774Yfs*5 promotes genomic instability through micronuclei formation"

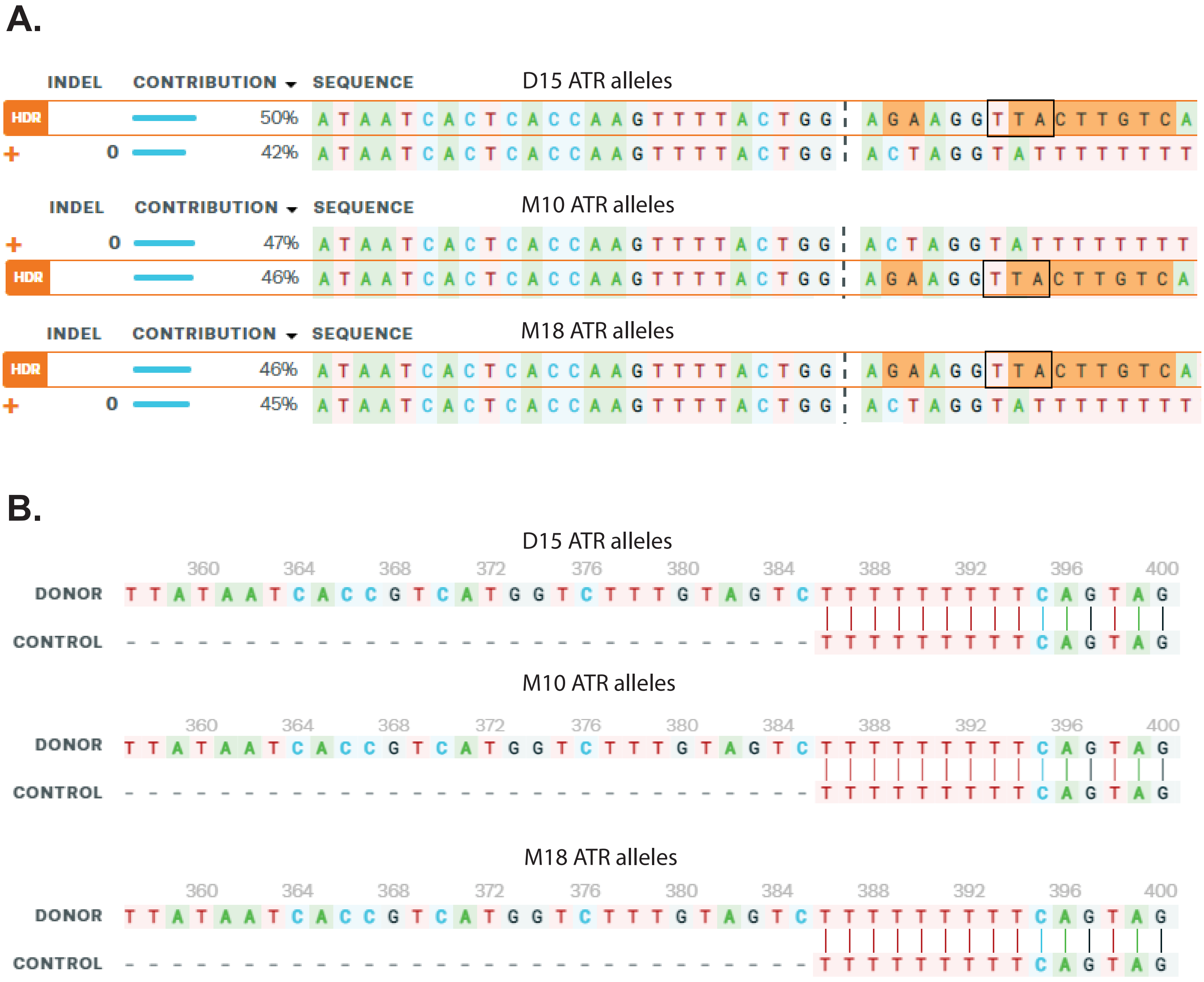
A.
INDEL
CONTRIBUTION
SEQUENCE
D15 ATR alleles
HDR 50% ATAATCACTCACCAAGTTTTACTGG AGAAGGTTACTTGTCA
+ 0 42% ATAATCACTCACCAAGTTTTACTGG ACTAGGTATTTTTTTT
M10 ATR alleles
+ 0 47% ATAATCACTCACCAAGTTTTACTGG ACTAGGTATTTTTTTT
HDR 46% ATAATCACTCACCAAGTTTTACTGG AGAAGGTTACTTGTCA
M18 ATR alleles
HDR 46% ATAATCACTCACCAAGTTTTACTGG AGAAGGTTACTTGTCA
+ 0 45% ATAATCACTCACCAAGTTTTACTGG ACTAGGTATTTTTTTT
B.
D15 ATR alleles
M10 ATR alleles
M18 ATR alleles
360 364 368 372 376 380 384 388 392 396 400
DONOR TTATAATCACCGTCATGGTCTTTGTAGTCTTTTTTTTTCAGTAG
CONTROL TTTTTTTTTCAGTAG

**Supplemental Figure 1: Validation of ATR heterozygosity in CRISPR knock-in cell lines. (A)** The relative proportion of each ATR allele in the three provided clones is shown, as determined by Inference of CRISPR edits (ICE) using Sangar sequencing. Observed alleles below 5% contribution are considered background noise and not included for analysis. Each clone (D15, M10, M18) shows two alleles at roughly 50% contribution apiece, one marked "+" for Wild-type, and the other marked "HDR" for the mutant allele. This allele contains a large insertion 5' of the cut site, where a CRISPR mediated knock-in of a 3x flag sequence followed by a premature stop codon occurred (Stop codon is shown in the black box). The sequences shown are for the template strand of ATR, so the stop codon (which is TAA on the coding strand) is observed 5' of the remaining insertion, part of which is shown in the alignment. **(B)** The alignment of the polynucleotide region of ATR (Poly-T on template strand) along with the beginning of the insertion of 3x flag in the three provided clones is shown, as determined by Inference of CRISPR edits (ICE) using Sangar sequencing.
Additional information about the sequence analyses can be found at
https://ice.synthego.com/#/analyze/results/44m837tq85q34twh/2066146_1_D15;F1
https://ice.synthego.com/#/analyze/results/antx6b8sfupg9ms7/2066146_1_M10;F1
https://ice.synthego.com/#/analyze/results/v6f8jmef4aae2ffx/2066146_1_M18;F1

Holcomb, et. al., Supplemental Fig. 2

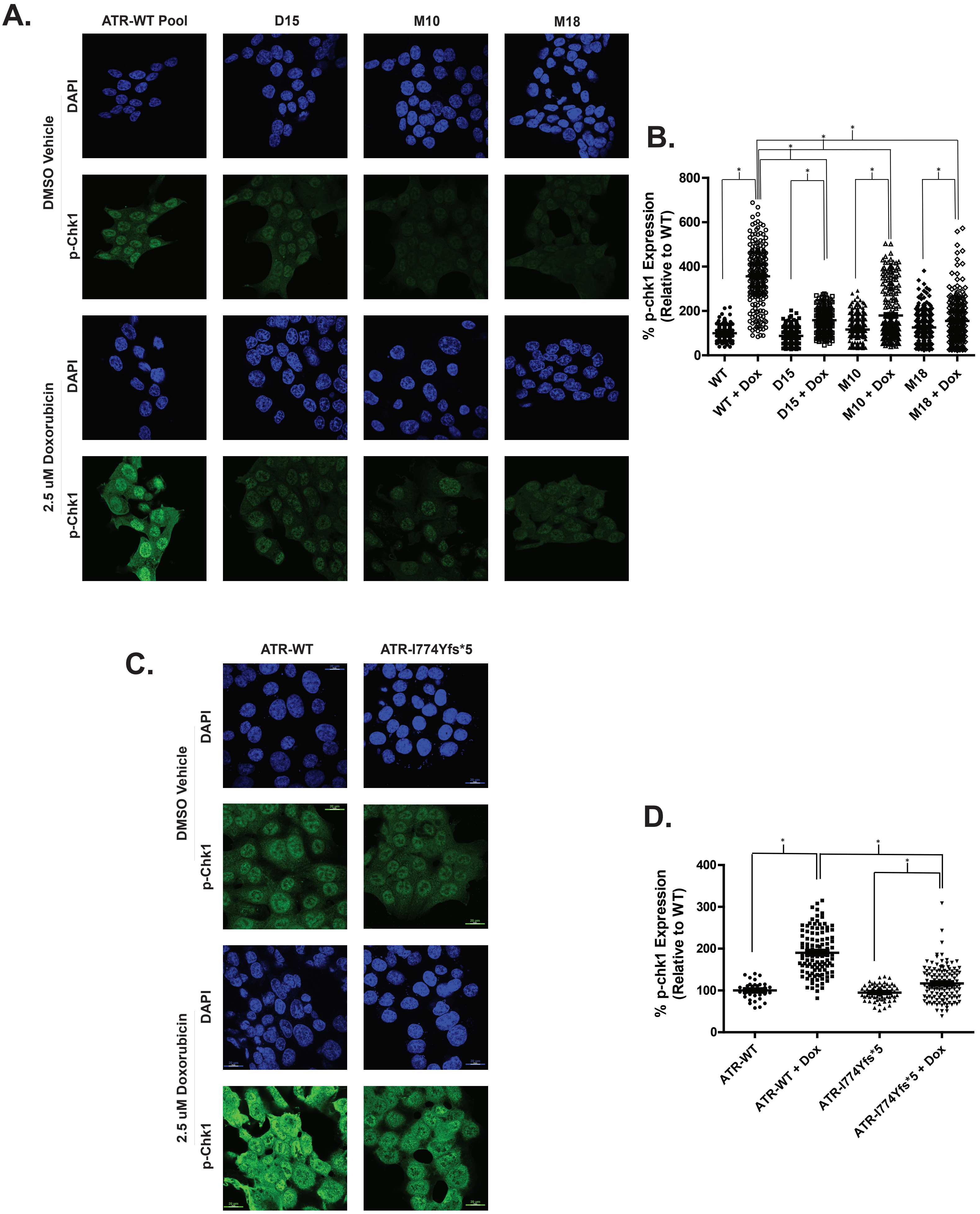

**Supplemental Figure 2: Confocal microscopy confirms ATR-I774Yfs*5 reduces doxorubicin mediated p-Chk1 induction in HCT-116 cells (A)** ATR-I774Yfs*5 heterozygotic mutant HCT-116 cell clones D15, M10, and M18 along with HCT-116 WT-ATR were seeded onto an 8-well chamber slide, grown overnight, and exposed to 2.5 µM Doxorubicin for 24 hours. After exposure, cells were harvested and analyzed by confocal microscopy. DAPI staining was used to define nuclear boundaries and p-Chk-1 (S317) was measured in each cell. (**B**) Densitometry analysis of pChk-1 levels across 3 independent experimental confocal replicates (mean ± standard deviation); minimum n of 200 cells per condition, * denotes statistical significance ($p < 0.05$) between groups compared (1-way ANOVA). **(C)** HCT-116 cells were seeded onto a chamber slide, transfected with ATR-WT or ATR-I774Yfs*5 for 24 hours and then exposed to 10 µM Doxorubicin or a vehicle control for 3 hours. After damage cells were immediately fixed, permeabilized, probed with p-Chk1 (S317) antibody and DAPI, and visualized by confocal microscopy. Nuclear p-chk1 intensity was determined on a per-cell basis and compared to the average value of the ATR-WT transfected cells. **(D)** A graphical representation of p-Chk1 (S317) expression in cells transfected with ATR-WT or ATR-I774Yfs*5 +/- doxorubicin treatment. Significance was calculated by a One-way ANOVA with a Bonferroni's Multiple Comparison Test and is denoted with asterisks between the indicated treatment conditions.

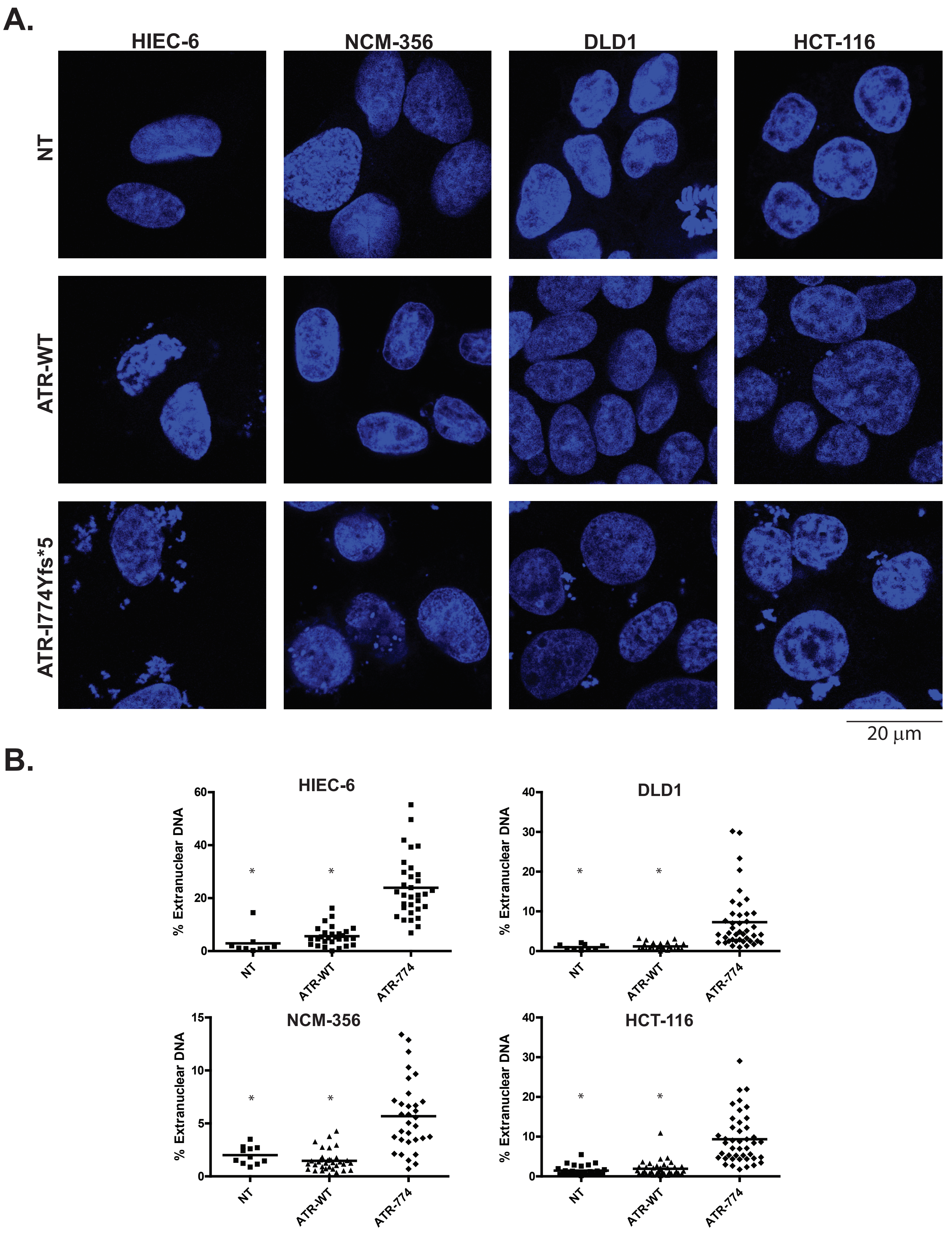
A.
HIEC-6
NCM-356
DLD1
HCT-116
NT
ATR-WT
ATR-I774Yfs*5
20 μm
B.
HIEC-6
DLD1
NCM-356
HCT-116
% Extranuclear DNA
NT
ATR-WT
ATR-774
*

**Supplemental Figure 3: Ectopic ATR-I774Yfs*5 generates a robust micronuclear phenotype (A)** HIEC-6, NCM-356, DLD-1, and HCT-116 colorectal cells were seeded into chamber slides and transfected with the indicated plasmids for 24 hours. After transfection, the cells were fixed, permeabilized, stained with DAPI for detection of DNA, and visualized by confocal microscopy. **(B)** Extranuclear DNA content was quantified from confocal images as a percentage of total DAPI content and plotted for each cell line across the transfection conditions; * denotes $p < 0.05$ comparing NT or ATR-WT to ATR-I774Yfs*5 using a 1-way ANOVA).

**A.**

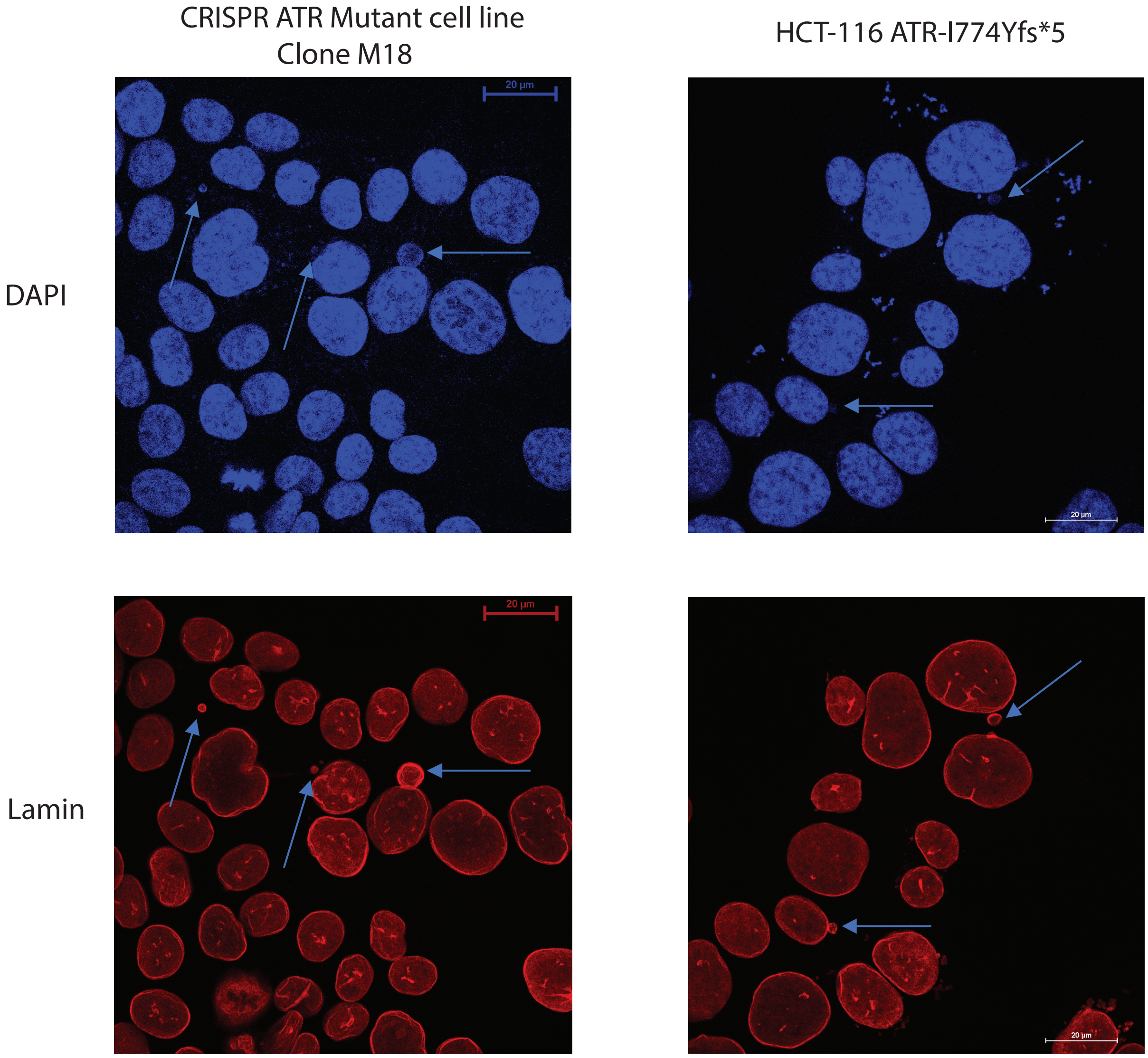


**B.**

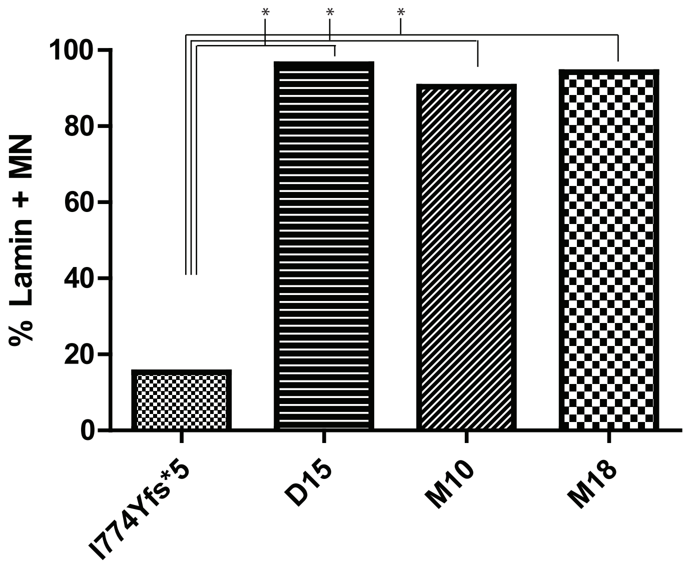


| | # MN | Lamin + MN | Lamin - MN | % Lamin + MN |
|---|---|---|---|---|
| ATR I774Yfs*5 | 59 | 9 | 50 | 15.3 |
| D15 | 54 | 52 | 2 | 96.3 |
| M10 | 94 | 85 | 9 | 90.4 |
| M18 | 52 | 49 | 3 | 94.2 |

**Supplemental Figure 4: MN produced by stable and transient ATR-I774Yfs*5 expression differ in lamin envelopment.** **(A)** Confocal images of HCT-116 cells with DAPI stain (above) and Lamin A/C (below) indicate that the majority (90-96%) of MN seen in the three stable, CRISPR-generated ATR mutant cells contain a lamin envelope (left panels), while transient transfection of HCT-116 cells with ATR-I774Yfs*5 (right panels) produce extranuclear DNA fragments that have a lower frequency (15%) of lamin coating. **(B)** A graph and table showing the incidence of lamin positivity in observed MN in the ATR-I774Yfs*5 transiently transfected or CRISPR-generated ATR I774Yfs*5 heterozygotic mutant cell lines; * denotes $p < 0.05$ comparing lamin frequency of MN between transient and stably expressing ATR mutant cell lines using Fisher's Exact test.
